# A Peek into the Plasmidome of Global Sewage

**DOI:** 10.1101/2021.03.08.434362

**Authors:** Philipp Kirstahler, Frederik Teudt, Saria Otani, Frank M. Aarestrup, Sünje Johanna Pamp

**Author notes:** Correspondence, Technical University of Denmark, 2800 Kongens Lyngby, Denmark.

## Abstract

Plasmids can provide a selective advantage for microorganisms to survive and adapt to new environmental conditions. Plasmid-encoded traits, such as antimicrobial resistance (AMR) or virulence, impact on the ecology and evolution of bacteria and can significantly influence the burden of infectious diseases. Insight about the identity and functions encoded on plasmids on the global scale are largely lacking. Here we investigate the plasmidome of 24 samples (22 countries, 5 continents) from the global sewage surveillance project. We obtained 105 Gbp Oxford Nanopore and 167 Gbp Illumina DNA sequences from plasmid DNA preparations and assembled 165,302 contigs (159,322 circular). Of these, 58,429 encoded for genes with plasmid-related and 11,222 with virus/phage-related proteins. About 90% of the circular DNA elements did not have any similarity to known plasmids. Those that exhibited similarity, had similarity to plasmids whose hosts were previously detected in these sewage samples (e.g. *Acinetobacter, Escherichia, Moraxella, Enterobacter, Bacteroides*, and *Klebsiella*). Some AMR classes were detected at a higher abundance in plasmidomes (e.g. macrolide-lincosamide-streptogramin B, macrolide, and quinolone), as compared to the respective complex sewage samples. In addition to AMR genes, a range of functions were encoded on the candidate plasmids, including plasmid replication and maintenance, mobilization, and conjugation. In summary, we describe a laboratory and bioinformatics workflow for the recovery of plasmids and other potential extrachromosomal DNA elements from complex microbiomes. Moreover, the obtained data could provide further valuable insight into the ecology and evolution of microbiomes, knowledge about AMR transmission, and the discovery of novel functions.

**Importance:** This is, to the best of our knowledge, the first study to investigate plasmidomes at a global scale using long read sequencing from complex untreated domestic sewage. Previous metagenomic surveys have detected AMR genes in a variety of environments, including sewage. However, it is unknown whether the AMR genes were encoded on the microbial chromosome or are located on extrachromosomal elements, such as plasmids. Using our approach, we recovered a large number of plasmids, of which most appear novel. We identified distinct AMR genes that were preferentially located on plasmids, potentially contributing to their transmissibility. Overall, plasmids are of great importance for the biology of microorganisms in their natural environments (free-living and host-associated), as well as molecular biology, and biotechnology. Plasmidome collections may therefore be valuable resources for the discovery of fundamental biological mechanisms and novel functions useful in a variety of contexts.

## Introduction

The term plasmid was introduced by Joshua Lederberg in 1952 to describe any extrachromosomal genetic particle (1). It was not until the 1970s when interest in plasmid research rapidly increased and plasmids were introduced as cloning vectors into an area that was dominated by phages as the vector for the transfer of pieces of DNA of choice (2). Since then, plasmids have been highly valuable tools in molecular microbiology. In their natural environment, plasmids are considered key players in horizonal gene transfer. They play crucial roles in the ecology and evolution of bacteria, including their pathogenicity as they can carry virulence factors such as toxins as well as antimicrobial resistances genes (3) (4–6). However, the global diversity of plasmids and distribution of antimicrobial resistance genes are yet to be revealed.

The presence of antimicrobial resistance genes on plasmids are of major interest in the clinical and veterinary areas since they can render prescribed antibiotics for treating pathogens ineffective. There have been a range of large-scale metagenomic-based surveys of antimicrobial resistance genes in soils, humans, animals, plants, and sewage (7–12). However, the genomic context of the AMR genes is largely unknown; for example, whether they are located in the bacterial genome or on plasmids. Such knowledge would be of great value to better assess their potential transmissibility rates and global impact of AMR-gene carrying plasmids on human health.

Plasmids are usually circular DNA elements in bacterial cells, but they can also occur in linear form and be present in archaea and eukaryotic organisms. The size of plasmids is highly variable, ranging from 1,000 bases to hundreds of kilobases. They are present in different quantities (copy numbers) in bacterial cells, varying from a single copy to hundreds of copies in a single cell. This intrinsic and unique nature of plasmids brings about several challenges in plasmidome research (i.e. research on the collective plasmid content in a sample). For example, the low plasmid/chromosome DNA ratio and potential low copy numbers can make it difficult to detect plasmids. These challenges are amplified when plasmidomes are examined from relatively low-cell-density environments such as wastewater. Even assembling and identifying plasmids with low copy number from high biomass samples including single isolates from whole genome sequencing (WGS) data can be challenging. To address these challenges, different approaches have been developed to increase the recovery of plasmids from WGS data (13–16).

Plasmids have now also been recovered from more complex microbiomes using a number of strategies. This includes multiple displacement amplification (MDA) with phi29 DNA polymerase prior to DNA sequencing (17), long read sequencing technology of plasmid DNA, or involvement of advanced assembly strategies (18–21). These studies have however been restricted to a single or few locations, and there is limited knowledge on similarity and differences between plasmids from a range of geographical locations (22–26). We recently showed differences in the AMR gene profiles in urban sewage around the globe, but the location of these AMR genes in the bacteria remains unknown (7).

To explore the plasmidome of global sewage, which is characterized by low bacterial cell numbers and challenges to isolate plasmid DNA as previously shown (23–27), we here employ an optimized plasmid DNA isolation procedure, followed by both, plasmid-safe DNase treatment and MDA to obtain sufficient plasmid DNA for Oxford Nanopore sequencing from global urban sewage samples. To improve plasmidome characterizations, we develop an assembly workflow, utilizing the long-read length from the Oxford Nanopore MinION sequencer and Illumina sequences. We obtain thousands of circular candidate plasmid sequences and explore their predicted function.

## Material and Methods

### Sample collection and preparation

From the global sewage sample collection (7), we selected 24 samples from 22 countries (see Table S1 in the supplementary material). The samples originated from the five most populated continents on Earth and for which we had sufficient sample material available. From each sample, a sewage pellet was collected from 250 ml untreated sewage by centrifugation at 10,000xg for 10 minutes at 5°C. The sewage pellets were stored at -80°C until use.

### Plasmid DNA extraction and enrichment

Plasmid DNA isolation was performed on individual sewage pellets (420 mg) using Plasmid Purification Mini Kit (Qiagen, Cat No./ID: 12123) with QIAGEN-tip 100 (Qiagen, Cat No./ID: 10043) following the manufacturer’s instruction with the following minor modifications: protein precipitation with P3 buffer mixture was incubated on ice for 15 minutes, elution buffer QF and EB buffer were preheated at 65°C prior to application, and the DNA pellet washing step was done using ice-cold 70% ethanol after isopropanol precipitation. LyseBlue dye for cell lysis indication was added, and all buffer volumes were adjusted to sewage pellet weight. The plasmid DNA pellet was dissolved in 35 μl EB buffer for 1 hour at room temperature. Linear chromosomal DNA was reduced by Plasmid-Safe ATP-Dependent DNase (Epicentre, USA) treatment for 24 hours at 37°C according to the manufacturer’s instructions. The DNase was inactivated at 70°C for 30 minutes. To selectively enrich for circular DNA, the plasmid-Safe DNase-treated DNA was amplified using phi29 DNA polymerase (New England Biolabs, USA) following the manufacturer’s instructions, similar to as previously described (22). The plasmid DNA is amplified through rolling circle amplification by the phi29 DNA polymerase using random primers, generating multiple DNA replication forks (17). This results in long DNA fragments that contain tandem copies (tandem repeats) of the same plasmid. Blank controls were used during plasmid DNA extractions and plasmid enrichment treatments. All negative controls had undetectable DNA measurements using Qubit double-stranded DNA (dsDNA) BR assay kit on a Qubit 2.0 fluorometer (Invitrogen, Carlsbad, CA).

### Plasmid DNA quality assessment

The plasmid DNA yields from the sewage samples were evaluated using gel electrophoresis and Qubit double-stranded DNA (dsDNA) BR assay kit on a Qubit 2.0 fluorometer (Invitrogen, Carlsbad, CA). Plasmid DNA purity was measured and validated by absorbance ratio of 260/280 and 260/230 using NanoDrop 100 (ThermoFisher). During pilot experiments that were aimed at protocol development and plasmid DNA enrichment, we also assessed the quality of our plasmid DNA preparations using a 2100 Bioanalyzer (Agilent).

### Library preparation and Oxford Nanopore sequencing

One μg plasmid DNA in 45 μl buffer was used for library preparation. DNA was used without fragmentation. End repair and dA-tailing were performed using NEBNext FFPE Repair Mix (New England BioLabs, 6630) and NEBNext® Ultra™ II End Repair/dA-Tailing Module (New England BioLabs, 7546). DNA was mixed with 3.5 μl NEBNext FFPE DNA Repair Buffer, 2 μl NEBNext FFPE DNA Repair Mix, 3.5 μl Ultra II End-prep reaction buffer and 3 μl Ultra II End-prep enzyme mix and volume was adjusted to 60 μl with nuclease-free water. The reaction tube was flicked 3 times and incubated at 20°C for 10 minutes, then inactivated by heating at 65°C for 10 minutes. Clean-up was done using 60 μl Agencourt AMPure XP beads. The beads-reaction suspension was incubated on a HulaMixer at the lowest speed for 10 minutes, followed by two washes with freshly prepared 70% ethanol. DNA was then eluted from the beads in 61 μl 65°C preheated nuclease-free water. A 1 μl DNA aliquot was assessed with Qubit dsDNA BR assay to ensure >700 ng were recovered. A volume of 60 μl of dA-tailed plasmid DNA were added to 25 μl Ligation Buffer (LNB), 10 μl NEBNext Quick T4 DNA Ligase NEBNext Quick Ligation Module (New England BioLabs, 6056) and 5 μl Adapter Mix (AMX), and mixed by flicking the tube 3-4 times followed by incubation at room temperature for an extended time of 1 hr. The adaptor-ligated plasmid DNA was cleaned up by adding 40 μl Agencourt AMPure XP beads, and the reaction was mixed by flicking the tube and followed by incubation on a HulaMixer at the lowest speed for 10 minutes. The beads were pelleted and resuspended twice in 250 μl Long Fragment Buffer LFB buffer (SQK-LSK109 kit, Oxford Nanopore Technologies). The cleaned adaptor-ligated DNA was then eluted by incubating the pellet in 15 μl Elution Buffer (SQK-LSK109 kit, Oxford Nanopore Technologies) for 20 minutes at room temperature, then transferring the supernatant to a new tube as constructed library. Flowcell priming and library loading preparation were performed according to the manufacturer’s instruction (SQK-LSK109 kit, Oxford Nanopore Technologies). Libraries were loaded on FLO-MIN106 R 9.4.1 Oxford Nanopore flowcells, and sequencing was run for 48 hours with MinKNOW software default settings.

### Illumina Sequencing

The enriched plasmid DNA samples were also subjected to Illumina NextSeq sequencing for downstream error-correction of contigs. Libraries were prepared using Nextera XT DNA Library Preparation Kit (Illumina, USA) following the manufacturer’s instructions. The libraries were sequenced using NextSeq 550 system (Illumina) with 2 × 150 bp paired-end sequencing per flow cell.

### Data processing

Basecalling of Nanopore reads was performed using the guppy basecaller (version 3.0.3+7e7b7d0) with the dna_r9.4.1_450bps_hac (high accuracy) configuration. Adapter trimming was performed using porechop (version 0.2.3) downloaded from https://github.com/rrwick/Porechop using the default parameters. Illumina sequencing data were quality and adapter trimmed using bbduk from the bbmap suite (https://sourceforge.net/projects/bbmap/, version 38.23) using the following settings: qin=auto, k=19, rref=adapters.txt, mink=11, qtrim=r, trimq=20, minlength=50, tbo, ziplevel=6, overwrite=t, statscolumns=5.

### Plasmid assembly from single Nanopore reads

Nanopore reads shorter than 10,000 bases were discarded. Each remaining read was cut into 1,500 bases long fragments and passed to the assembly step. The initial fragmentation step of the reads is needed since each read, amplified from a circular element during sample preparation, consists of multiple tandem repeats of the circular element. This is done to eliminate the tandem repeats as well as increase the accuracy of the resulting candidate plasmid DNA sequence. We set the cutting threshold to 1.5 kb to balance between preserving the benefits of long read sequencing and accounting for the error rate of Nanopore sequencing. We decided for a length threshold for cutting (i.e. 1.5 kbp) to not create candidate plasmid DNA sequences from small plasmids that contain multiple copies of the same plasmid. We set the cutting threshold to 1.5 kbp to balance between preserving the benefits of long read sequencing and the error rate of Nanopore sequencing. We also preferred to keeping the cutting threshold more towards the short range to not create candidate plasmids form small plasmids that contain multiple copies of the same plasmid sequence. Read fragments originating from one single read were assembled using minimap2 (version 2.17-r943-dirty) in combination with miniasm version 0.3-r179 (parameter -s 800bp), and error corrected using racon version 1.3.3 (28–30). Assembled contigs were discarded if, after mapping the assembled contig back to the original Nanopore read, hits did not span more than 60% of the read, and if two hits overlapped by more than 50 bp. Assembled candidate contigs were error-corrected using 5 iterations of pilon version 1.23 using the respective Illumina reads from the same sample (31). Candidate contigs longer than 1,000 bases were used for downstream analyses. A schematic overview of the method is presented in Figure 1A.

**Figure 1.**
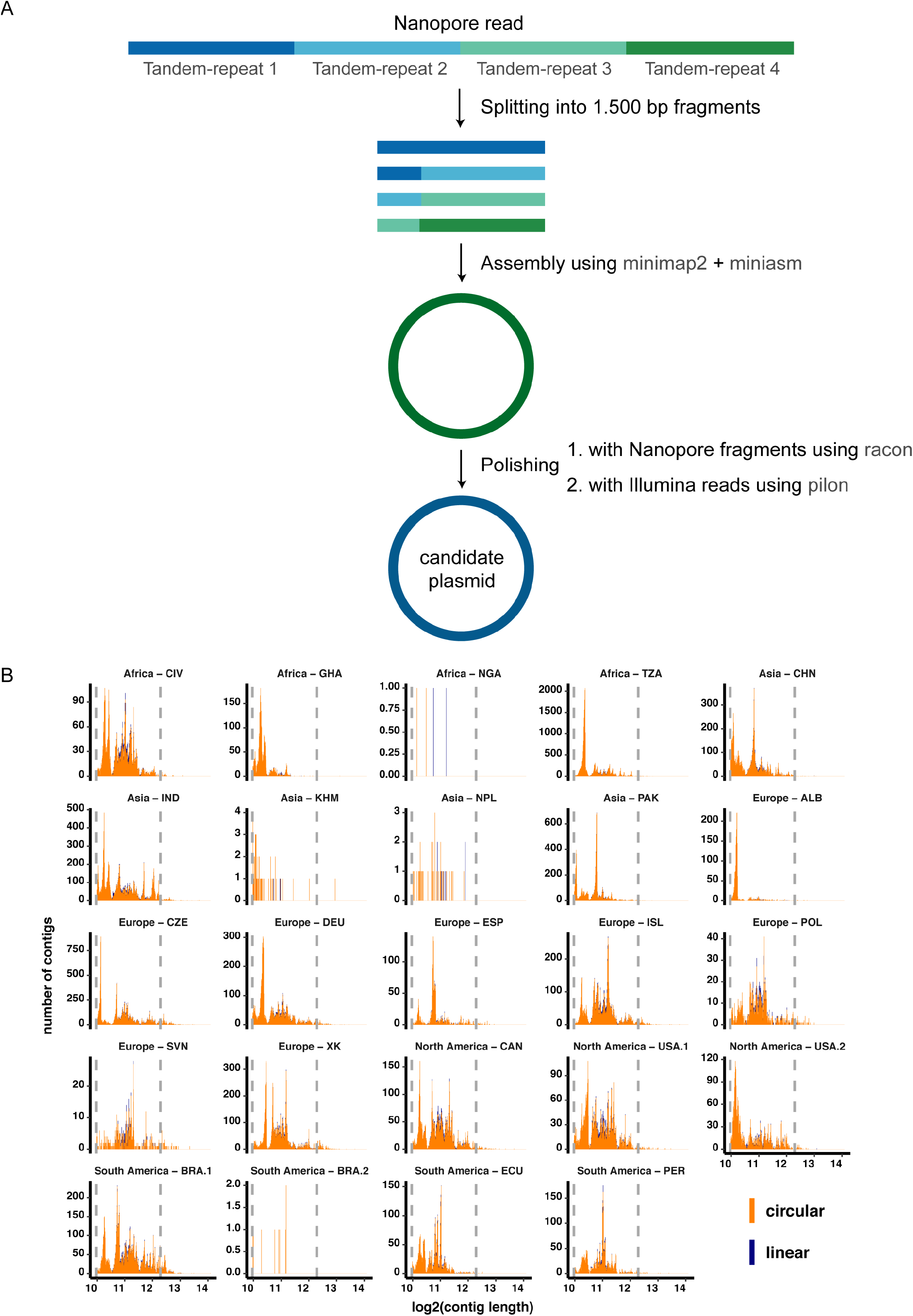
Schematic overview of the single read assembly workflow and size distribution of assembled reads. A) Nanopore reads (based on plasmid DNA amplified with phi29) longer than 10,000 bases were split into 1,500 bases long fragments. The sequence fragments were then assembled using minimap2 and miniasm and subsequently polished two times: 1. with the Nanopore fragments using racon and 2. with the Illumina reads using pilon. B) The size distribution of circular (orange) and linear (violet) assembled elements. These are the candidate plasmid sequences that successfully mapped to the original Nanopore read (i.e. covering more than 60% of the read, and not overlapping by more than 50 bp for multiple hits). Of the total 165,302 assemblies, 159,322 were characterized to be circular and 5,980 to be linear.

### Global plasmidome analysis

To examine the obtained plasmids from our global sewage collection in relation to already known plasmids, we compared our obtained candidate plasmid DNA sequences to the DNA sequences in the plasmid database (PLSDB) using the webtool of PLSDB version 2019_10_07 (32). We used search strategy ‘Mash screen’ with a maximum p-value of 0.005 and minimum identity of 95%, as well as the option ‘winner-takes-all strategy. Samples with less than 100 circular assembled contigs were removed from the analysis as well as genera with less than 10 occurrences over all samples. A clustering of samples was performed using Euclidean distance of the clr-transformed values. Furthermore, all candidate plasmid sequences were sketched using MASH version 2.2 (33). The MASH-distances between all samples were calculated using default settings, resulting in a 24 by 24 distance table that was used for principal component analysis.

### Antimicrobial resistance gene detection analysis

The trimmed Nanopore and Illumina reads were mapped against the ResFinder database (2020-01-25) using kma (version 1.3.0) (34, 35). The Nanopore reads were mapped with settings: mem_mode, ef, nf, bcNano, and bc=0.7. Illumina reads were mapped with settings: mem_mode, ef, nf, 1t1, cge, and t=1. Resistance genes were counted across variants, for example the alleles tet(A)_4_AJ517790 and tet(A)_6_AF534183 were both counted as tet(A). Centered log ratios were calculated using the pyCoDa package (https://bitbucket.org/genomicepidemiology/pycoda/src/master/).

### Gene prediction and functional analysis

Gene prediction was performed using prodigal version 2.6.3, and annotation of protein families was done using hmmscan from HMMER3 version 3.3.1 (http://hmmer.org/) against the pfam database version 33 (36, 37). Predicted genes as well as functional annotation were rejected if the p-value was above 0.000001. Gene ontology (GO) annotations for Pfam IDs were acquired using the mapping of Pfam entries to GO terms as described by Mitchell *et al*. (38).

To distinguish between potential plasmid and non-plasmid contigs, we used a scheme described previously (39). The scheme contains Pfam identifiers highly specific for plasmids and viruses. Proteins with a plasmid replication initiator protein Rep_3 (PF01051) domain (n=24,824) were investigated further together with the full set of reference Rep_3-domain proteins (n=1,637) downloaded from Pfam (version 33.1). The two data sets were combined and Rep_3-domain proteins with a length of <40 aa residues were discarded, resulting in a data set of 16,930 Rep_3 (PF01051) domain proteins. The protein sequences were aligned using MAFFT (version 7.221) as part of the Galaxy platform (40, 41). A phylogenetic tree was then build using FastTree (version 2.1.10) (42) and visualized using FigTree (version 1.4.4) (https://github.com/rambaut/figtree/releases).

### Generation of plasmid maps

The 50 longest assemblies from each sample were annotated using Prokka (43). Contigs of interest were chosen for mapping based on the presence of known plasmid-encoded genes, such as replication and mobilization systems, toxin-antitoxin pairs, and AMR genes. Plasmids were inspected and visualized using DNAPlotter (44) and Geneious Prime version 2020.2.4 (www.geneious.com). If a coding sequence (CDS) from the Prokka analysis remained unannotated, it was manually annotated by using BLAST search function against the nr database (45) and scanned with HMMER3 against the Pfam database as described above.

## Results

### Nanopore and Illumina sequencing ouput from plasmid DNA-enriched global sewage samples

The sequencing of 24 plasmid-enriched DNA preparations from untreated sewage from 5 continents (Africa, Asia, Europe, North America, and South America) using Oxford Nanopore sequencing technology produced 1.2 to 9.7 Gbp (median 3.5 Gbp) sequencing data per sample (see Table S1 in the supplementary material). The median read length was 7.3 kb (range 1,075 to 11,018 bases) (see Figure S1 in the supplemental material). After quality trimming and removing sequences below 10,000 bases, the median sequencing throughput was 1.9 Gbp and the median read length 23,000 bases (see Table A at https://doi.org/10.6084/m9.figshare.13395446). The Illumina generated sequencing data per sample were between 1.5 and 9.7 Gbp with a median of 4.8 Gbp after adapter and quality trimming. A median of 41 million paired-end reads per sample were obtained (see Table B at https://doi.org/10.6084/m9.figshare.13395446).

### Circular DNA sequences obtained using single Oxford Nanopore reads

Upon assembly and polishing (Figure 1A), we obtained a total of 165,302 contigs from the 24 sewage samples, of which 159,322 contigs (96.4%) were suggested by miniasm to be circular (Figure 1B, and see Table C at https://doi.org/10.6084/m9.figshare.13395446). The longest assembled circular contig had a length of 17.4 kbp and was obtained from a sample in Brasil (BRA.1, South America). Most of the circular contigs were obtained from the Tanzanian (TZA, Africa) sewage sample, and they had an average length of 1.7 kbp (see Table C at https://doi.org/10.6084/m9.figshare.13395446).

### Classification of assembled circular DNA elements

To obtain information about the identity of the obtained circular DNA elements, we performed gene prediction, annotation, and classification based on plasmid- and virus/phage-specific Pfam domains (39). Overall, we detected Pfam domains (including domains of unknown function (DUF)) on 47.01% of the circular elements, potentially suggesting the presence of many novel DNA sequences not encoding for known protein domains. For the DNA elements (circular & linear) for which Pfam domains were detected, the majority (88.39%) contained predicted genes with plasmid- or virus/phage-related Pfam entries (see Figure 2, Figure S2 in the supplementary material, and Table D at https://doi.org/10.6084/m9.figshare.13395446). Overall, we found 55,337 circular DNA elements that encoded for known plasmid-related Pfam domains (and not viral-related Pfam domains). The highest number of plasmid-related candidate sequences were detected in the sample from the Czech

**Figure 2.**
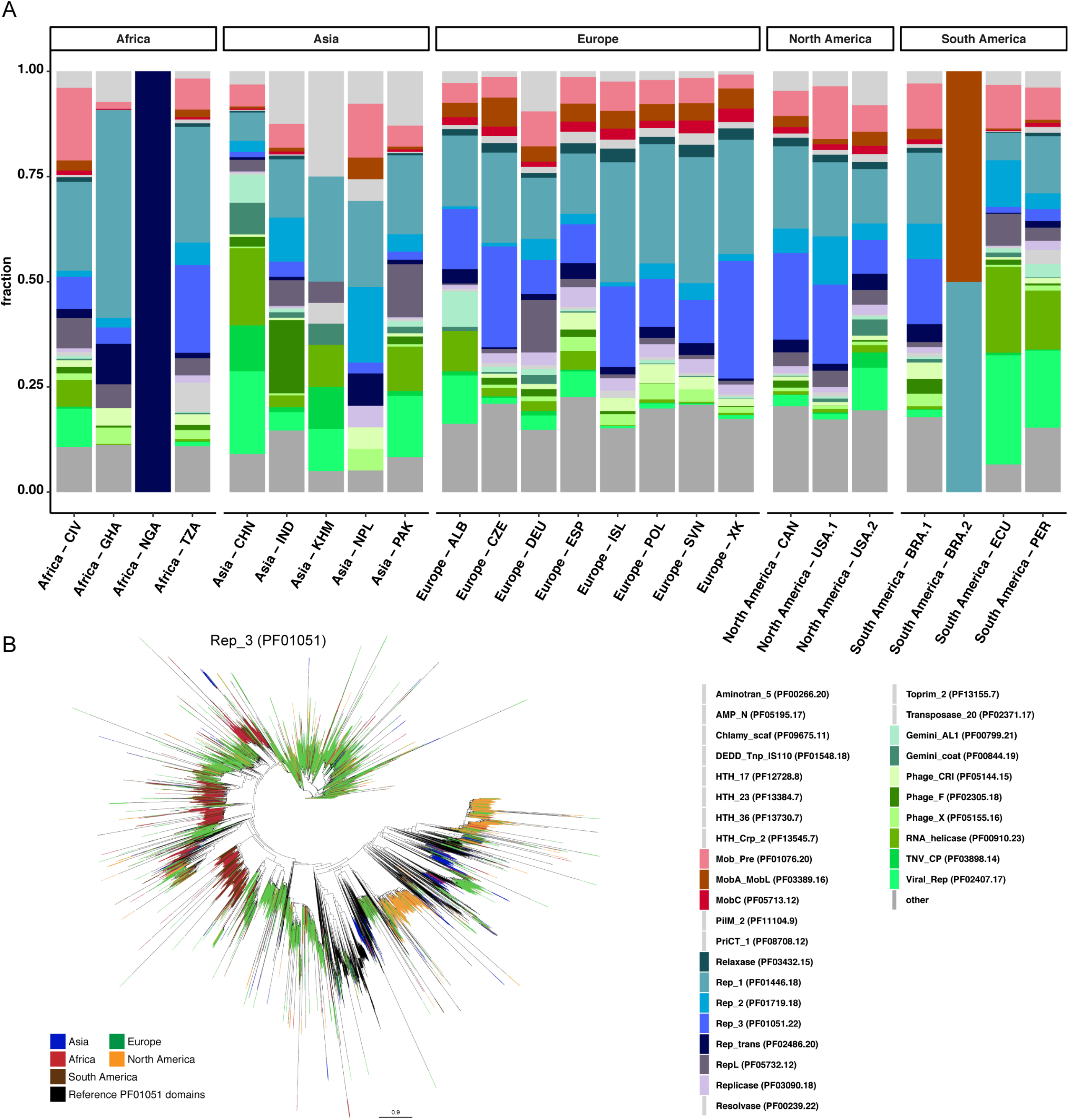
Functional characterization of circular DNA elements based on protein families. A) The bar plot displays the fraction of Pfam identifiers assigned to predicted proteins on the circular elements. The 31 Pfam identifiers represent the Top10 Pfam identifiers for each sample. Protein domains specifically involved in plasmid mobilization and plasmid replication are indicated by red and blue colors, respectively (see legend to the bottom right). Virus/phage related Pfam identifiers are indicated in green colors. Remaining Pfam identifiers are grouped (other) and indicated by dark grey. B) The dataset of proteins with a Rep_3 (PF01051) domain (n= 24,824) were combined together with the 1,637 reference Rep_3 (PF01051) proteins from Pfam. The protein sequences with a length of >/= 40 aa (n=16,930) were aligned using MAFFT. A phylogenetic tree was build using FastTree and visualized using FigTree. A high-resolution version of the phylogenetic tree is available from Figshare at https://doi.org/10.6084/m9.figshare.14112992.

Republic (CZE, Europe), followed by Tanzania (TZA, Africa), and Kosovo (XK, Europe). The sample from China (CHN, Asia) was the only sample from which more potential virus/phage-related contigs than candidate plasmids were obtained (see Figure 2, Figure S2 in the supplementary material, and Table D at https://doi.org/10.6084/m9.figshare.13395446).

On the circular elements with plasmid-related Pfam domains, protein families involved in plasmid replication were the most abundant and they included Relaxase, *Rep_1, Rep_2, Rep_3, Rep_trans, RepL, and Replicase* (Figure 2A). For example, we detected a total of 24,824 open reading frames with a plasmid replication initiator protein Rep_3 (PF01051) domain. Even though Rep_3-domain proteins from all continents were observed across the phylogenetic tree, some clades mainly represented proteins from one continent, interspersed with protein sequences from other continents (Figure 2B). For instance, clades that mainly harbored proteins originating from Europe, also frequently contained protein sequences from North America and other continents. Clades dominated by Rep_3 (PF01051) domain proteins from Africa also frequently harbored similar proteins from South America.

Furthermore, protein families involved in plasmid mobilization were detected, such as Mob_Pre, *MobA_MobL*, and *MobC* (Figure 2A). In addition, we identified protein families related to virus/phage replication and capsid proteins, as well as protein domains binding to DNA (HTH_17, HTH_23, HTH_Crp_2) and that might be involved in regulating gene expression.

### Global plasmidome pattern based on known plasmids

To examine whether our collection of plasmid sequences contained already known sequences, we compared the obtained plasmid DNA sequences to the entries in the plasmid database (PLSDB). This analysis revealed that only 10.1% of our circular elements were similar to known plasmids (see Table E at https://doi.org/10.6084/m9.figshare.13395446). The majority of plasmids that exhibited some similarity to entries in the PLDB originated from *Acinetobacter* (33%), *Enterococcus* (21%) as well as *Flavobacterium* (10%); genera that were previously detected in these sewage microbiomes (7).

Overall, most plasmids with similarities to already known ones were found in the samples from India, Kosovo, Pakistan, Czech Republic, Iceland, and Brazil (see Table E at https://doi.org/10.6084/m9.figshare.13395446). Clustering analysis of the abundancies of plasmids with known relatives in PLSDB revealed three main clusters (Figure 3A). The first cluster comprised samples that overall exhibited a low number of known plasmids and included samples from Europe (ALB, POL, ESP, SVN) and a sample from Ghana. The second cluster included samples with plasmids from a large range of bacterial genera at higher abundance, and comprised samples from Europe (ISL, DEU, CZE), North America (USA.1, USA.2, CAN), India, Brazil and Tanzania. The third cluster comprised samples with known plasmids from few bacterial genera and included samples from Asia (CHN, PAK), Africa (CIV), Europe (XK), and South America (ECU, PER) (Figure 3A).

**Figure 3.**
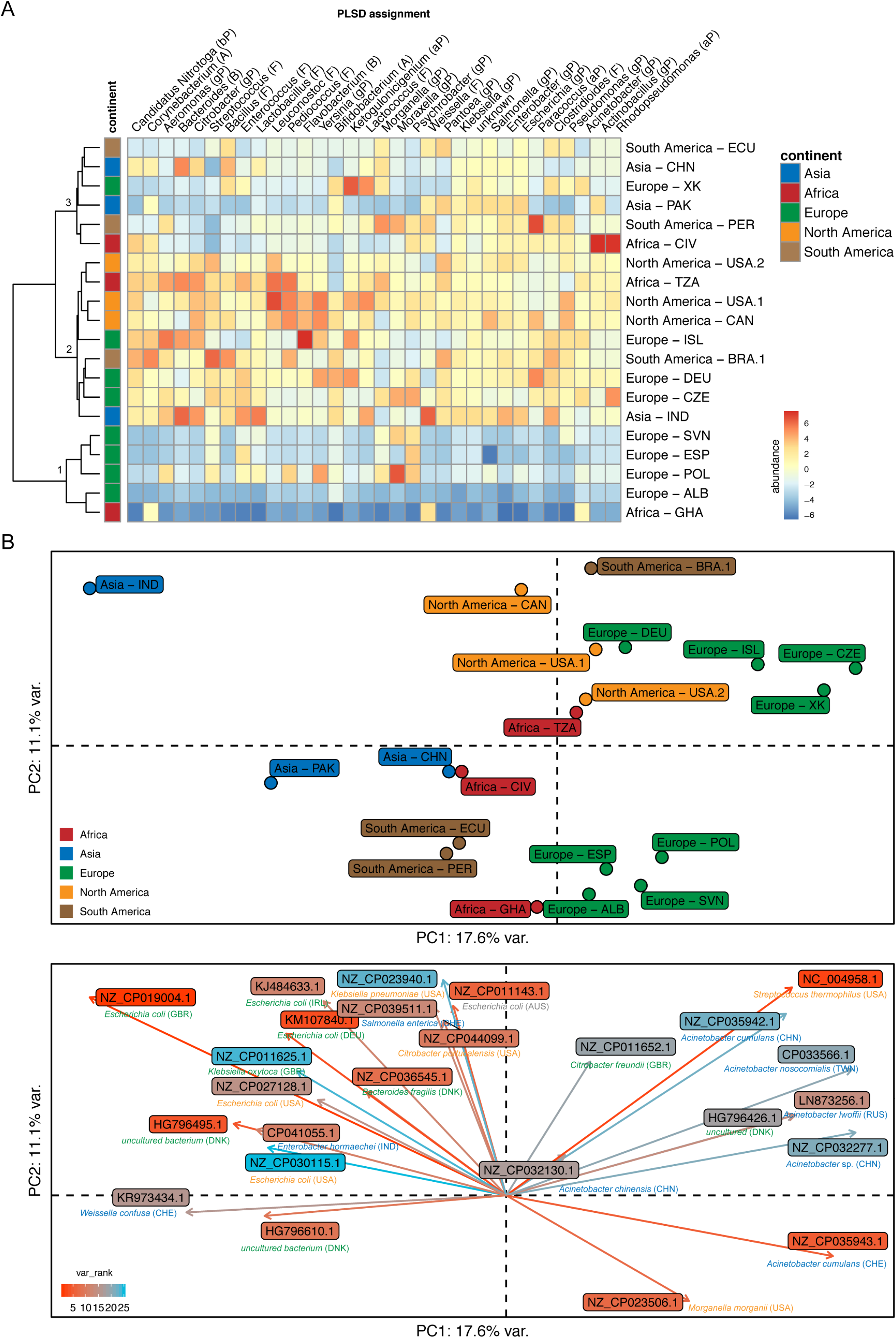
Comparison of candidate plasmids from global sewage with known plasmids in plasmid database (PLSDB). A) Heat map of centered log ratio (clr)-transformed abundancies of plasmid candidates assigned to plasmids in the PLSDB at bacterial genus level. The phylum level is indicated in parenthesis, A: Actinobacteria; B: Bacteroidetes; aP: alpha-Proteobacteria; bP: beta-Proteobacteria; gP: gamma-Proteobacteria; F: Firmicutes. Clustering of samples was performed using Euclidean distance of the clr-transformed values. B) Principal component analysis of clr-transformed abundancies of known plasmids detected by the PLSDB. The plot on the top reveals similarities and differences between samples. The plot in the bottom reveals the known plasmids that drive the partitioning of the samples, with 17.6% of the variation explained by the first and 11.1% by the second principal component.

In a principal component analysis of the same data, a similar clustering was observed. Furthermore, along the first principal component, samples from Asia and Europe appeared to be most different from each other and with samples from Africa, and North and South America in between. Upon examining the particular reference plasmids and their bacterial hosts that were driving this pattern a similar observation was made: plasmids from bacterial hosts originating from Europe appeared to segregate along the first principle component from plasmids and their bacterial hosts originating from Asia (Figure 3B). This observation was supported by a cluster analysis on plasmid-level, in which five clusters were observed: Samples from Europe did not cluster with samples from Asia, and different sets of known plasmids were found in the samples from Europe and Asia, respectively (see Figure S3 in the supplemental material). Generally, only few known plasmids were detected in the samples from Albania, Slovenia, Spain, Poland, Ecuador, and Ghana (see Figure S3 in the supplementary material and Table E at https://doi.org/10.6084/m9.figshare.13395446).

Given the large fraction of candidate plasmid sequences that did not exhibit similarity to already known plasmids, we performed a reference-independent analysis by calculating Mash-distances based on all plasmid sequences for each sample. In this analysis, the plasmidomes clustered in two main clusters (see Figure S4 in the supplemental material). The first cluster harbored all samples from Europe (with the exception of Poland), as well as the samples from Canada (North America), Pakistan and India (Asia), and Côte d’Ivoire (Africa). The second cluster harbored all samples from South America, both samples from the USA (North America), as well as Tanzania and Ghana (Africa), and China (Asia) (see Figure S4 in the supplemental material). This suggests that the sequence space encompassing novel plasmid sequences (i.e. those that did not exhibit similarity to sequences in the PLSDB) provides an extended, yet to be discovered, dimension into plasmid ecology and evolution.

### Antimicrobial resistance genes in plasmidomes

To gain insight into antimicrobial resistance genes on the plasmids from sewage, and compare them to those detected in the whole community of the same sewage samples, we performed a ResFinder analysis on three sequencing read data sets: whole community DNA sequenced using Illumina (7), plasmidome DNA sequenced using Illumina (this study), and plasmidome DNA sequenced using Nanopore sequencing (this study).

Overall, many of the antimicrobial resistance genes and antimicrobial classes that were detected using whole community sequencing, were also detected in the two plasmidome datasets, with a few exceptions. For example, the two antimicrobial classes macrolide-streptogramin B and lincosamide-pleuromutilin-streptogramin A were not detected in the plasmidome samples in about half of the cases (Figure 4A, and see Figure S5A in the supplementary material, and Tables F and G at https://doi.org/10.6084/m9.figshare.13395446). Occasionally, also genes conferring resistance to other antimicrobial classes were not detected in individual plasmidome samples as compared to the whole community, and these included genes conferring resistance to lincosamide, phenicol, or aminoglycoside. It may be that genes that were detected more frequently in the whole community sample, as compared to the plasmidome samples, are preferentially encoded on the bacterial chromosomes or larger plasmids.

**Figure 4.**
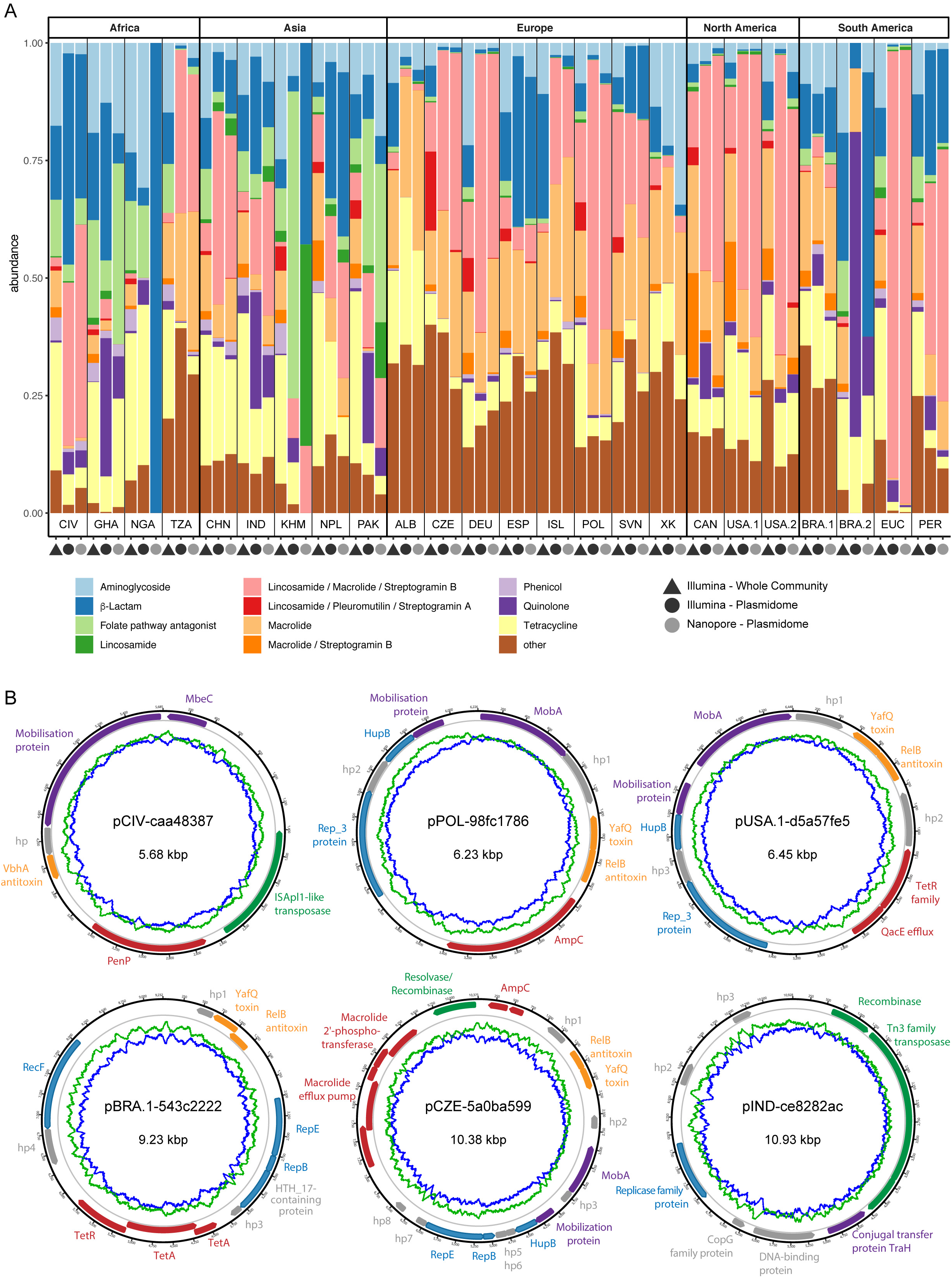
Antimicrobial resistance profiles from the whole community and plasmidomes from global sewage. **A)** Bar plot displaying the proportions of antimicrobial resistance classes detected in a ResFinder-based analysis using the Illumina reads from the whole community, as well as Illumina reads from the plasmid preparations and Nanopore reads from the plasmid preparations. B) Six examples of candidate plasmids are visualized in plasmid maps. The outermost black circle indicates the plasmid chromosome, the coding sequence regions are colored according to their predicted function: replication (blue), mobilization (violet), transposition of DNA (green), antimicrobial resistance (red), toxin-antitoxin systems (orange), hypothetical proteins (hp) and other proteins (grey). The blue and green line indicate the GC and AT-content, respectively. The plasmids are named according to their origin, CIV (Côte d’Ivoire), POL (Poland), USA.1 (USA), BRA (Brasil), CZE (Czechia), and IND (India). Some sequencing errors might still be present in the candidate plasmid sequences, which are likely the reason why a few open reading frames are not properly predicted and appear fragmented, such as the gene encoding for AmpC and Macrolide efflux pump genes in the plasmid from Czechia. A detailed description about the plasmids is available from Figshare at https://doi.org/10.6084/m9.figshare.14039390.

Conversely, genes conferring resistance to the antimicrobial classes macrolide-lincosamide-streptogramin B, as well as macrolide, and quinolone, were more frequently observed in the plasmidome samples (Figure 4A, and see Figure S5 in the supplementary material and Tables F and G at https://doi.org/10.6084/m9.figshare.13395446). The most frequently observed AMR genes related to these three classes were *ermB, ermT, ermF* (macrolide-lincosamide-streptogramin B), *mphE, mefA, msrD* (Macrolide), and *qnrB19, qnrD1, qnrD2, qnrD3, qnrVC4* (Quinolone). The higher frequency of those genes in the plasmidome samples may suggest that they are more frequently found on plasmids in general, or on smaller plasmids as compared to large ones. Another gene that was frequently observed across samples is *msrE*, and which was slightly higher abundant in plasmidomes (average abundance 15.4%, SEM 1.86) as compared to whole community samples (average abundance 11.5%, SEM 1.88). As examples, a few randomly chosen candidate plasmids and their encoded genes, including AMR genes, are displayed in Figure 4B.

### Functional characterization of plasmidomes

To gain further insight into the functions encoded on all circular elements, we obtained GO annotations for the predicted proteins through mapping of pfam entries to GO terms. A clustering analysis revealed the separation of plasmidomes into two main clusters (see Figure S6 in the supplemental material). Cluster 1 comprised samples from Europe (ISL, CZE, XK, DEU) as well as North America (USA.1, CAN) and South America (BRA.1, ECU). Cluster 2 comprised the samples from Asia (IND, PAK, CHN), Africa (TZA, CIV) and the remaining samples from Europe (POL, ESP, SVN) and South America (PER). This clustering based on protein functions appeared to have some similarity to the clustering based on nucleotide sequence similarity to known plasmids (Figure 3). In both analyses, the European samples from ISL, CZE, and DEU exhibited similarities, while the other European samples from POL, ESP, SVN clustered together separately. Furthermore, in both analyses, samples from North America (USA.1, CAN) and South America (BRA.1) clustered with the European samples from ISL, CZE, and DEU.

Functions that appeared to be enriched in samples from cluster 1 include, conjugation, recombinase activity, DNA methylation, protein secretion (type IV secretion system), response to antibiotic, toxic substance binding, response to toxic substance, unidirectional conjugation, and bacteriocin immunity (see Figure S5 in the supplemental material). Cluster 2 appeared to overall have fewer proteins that could be annotated using this strategy, and the samples exhibited a higher diversity of functional patterns compared to samples from cluster 1. Some samples from cluster 2 exhibited an enrichment of proteins that may be related to viruses/phages, such as viral capsids, structural molecule activity, RNA binding, RNA helicase activity, and these were in particular samples that appeared to have a higher abundance of virus/phage related Pfam domains (Figure 2). The majority of samples in both clusters harbored proteins involved in plasmid maintenance (see Figure S6 in the supplemental material).

## Discussion

This is the first study to investigate plasmidomes at a global scale using long read sequencing from sewage. We show that our approach facilitated the recovery of complete plasmids from complex metagenomic samples with a sufficient quality to perform gene prediction and functional annotation. In total, we obtained 165,302 DNA elements of which 159,322 were circular. The average length was 1.9 kb (min 1 kbp, max 17.4 kbp), suggesting that mainly small plasmids were obtained. This might reflect the true distribution but could also be biased due to a number of reasons, for example, smaller plasmids are more stable and thus have higher chance of getting though the DNA extraction step undamaged. Since a DNase step was used to reduce the amount of chromosomal DNA, damaged plasmids might have been digested as well. Another possibility could be that some plasmids were already damaged during storage and transportation, as the sewage was frozen and shipped, and many of the samples arrived thawed and were frozen again. Another reason could be that our assembly workflow was not able to perform a successful assembly on larger plasmids with a high number of tandem-repeats.

We identified a range of functions encoded on the candidate plasmids, including plasmid replication and maintenance, mobilization, conjugation, antimicrobial resistance, and bacteriocin immunity. However, not all plasmid-related DNA elements encoded for a plasmid-replication gene, suggesting that they may not be self-replicating DNA molecules. It should though be noted that also already described plasmids do not necessarily encode for a rep gene using current annotation algorithms. Furthermore, we found that about half of the circular DNA elements did not encode for any known Pfam domains. This could suggest that we detected many novel DNA sequences not encoding for known protein domains. A hypothesis could be that a fraction of the circular DNA elements are novel extrachromosomal elements that are hitherto undescribed and may also originate from various domains of life, including bacteria, archaea, and eukaryotes (46–48). Alternatively, open reading frames might not always have been properly detected because of sequencing errors not corrected in the polishing steps with Nanopore and Illumina reads. This could certainly have contributed to it, as we occasionally observed fragmented genes due to remaining sequencing errors, even after polishing. This challenge may be alleviated with the ongoing improvement of Oxford Nanopore chemistry and basecalling algorithms. Nevertheless, collectively, we obtained 58,429 DNA elements (circular & linear) that encoded for proteins with plasmid-related Pfams, and 17,292 circular DNA elements exhibited sequence similarity to known plasmids, suggesting that we successfully discovered many novel candidate plasmid DNA sequences.

For candidate plasmids that exhibited some similarities to known plasmids, we found that they originated from bacterial taxa previously detected in these complex sewage samples, such as *Acinetobacter, Escherichia, Moraxella, Enterobacter, Bacteroides*, and *Klebsiella* (7). These genera include bacteria that are part of the human gut microbiome and/or opportunistic pathogens. Hence, some of these plasmids might play a role in gut microbial ecology and potential antimicrobial resistance transmission (49, 50). It should be noted, however, that overall, only ∼10.1% of our circular elements were similar to known plasmids in the PLSDB, and which may be partly explained by differences in plasmid contents (plasmid average size 1.9 kbp (this study) and 53.2 kbp (PLSDB)) (32). In line with this, we observed that the plasmidome samples clustered somewhat differently when all candidate plasmid sequences were taken into account (and not only those that exhibited similarity to known reference plasmids). It will be interesting to investigate our candidate plasmids further in future studies, ideally through involvement of more plasmidome samples and extended metadata. There may be a range of factors that may play role in explaining differences and similarities between plasmidomes, such as climate, population-related differences including human ethnicity, health status, sanitation, and economy including trading between countries.

Overall, AMR classes that were detected in the plasmidome sequencing data sets were also found in the sequencing data from the whole complex sewage samples, suggesting that the plasmidomes are a good representation of what is present in the complex samples. Some AMR gene classes, however, were more predominant in the whole community (e.g. macrolide-streptogramin B, lincosamide-pleuromutilin-streptogramin A), and others more in the plasmidomes (e.g. macrolide-lincosamide-streptogramin B, macrolide, and quinolone). This could suggest that the AMR genes conferring resistance to the latter AMR gene classes are preferentially located on plasmids as compared to chromosomes. However, given that we mainly recovered small plasmids, it could also be an indication for that the AMR genes preferentially detected in the whole community may be located on large plasmids that were not recovered here. Whether certain abundant AMR genes in the plasmidomes are plasmid- or chromosome-associated may also be dependent on the particular bacterial host (see Figure S7 in the supplemental material) (51).

While our approach and findings are a significant advancement to previous work, there are still aspects that can be improved in the future. For example, the assembly workflow could be improved to resolve remaining repetitive regions within the plasmid, as a range of circular elements still consisted of tandem-repeats of the actual plasmid sequence. This could potently be solved by introducing a dynamic cutting step using the k-mer composition of the full read. Despite the high error rate of the Nanopore sequencing reads, the raw read should still contain a set of k-mers with 10-15 bases length that could help interfering the appropriate fragmentation length. In addition, the plasmid DNA isolation could be improved significantly to increase a) the overall amount of plasmid DNA (in order to avoid having to perform MDA), and b) the amount of larger plasmids. Further possibilities to identify new plasmids could also involve *in vivo* proximity-ligation Hi-C or single-cell sequencing that would also allow the discovery of new plasmids directly together with their host cell (52, 53).

Overall, our study provides new insight about the technical applicability of long-read Nanopore sequencing for plasmidome analysis of complex biological samples, as well as a foundation for exploring plasmid ecology and evolution at a global scale. For example, we can now better explore the genomic context of AMR genes, and reveal whether they are located on the microbial chromosome or on mobile genetic elements such as plasmids. This knowledge is of great value in assessing the potential transmissibility of AMR genes with resulting impact on antibiotic treatments in the medical and veterinary sectors and the one health perspective. Furthermore, the dataset provides a valuable resource for further exploring extrachromosomal DNA elements including potential novel functions.

## Acknowledgment

This work was mainly supported by The Novo Nordisk Foundation (NNF16OC0021856: Global Surveillance of Antimicrobial Resistance), and partially by the European Union’s Horizon 2020 Research and Innovation Programme under grant agreement No 773830: One Health European Joint

Programme. The funders had no role in study design, data collection and interpretation, or the decision to submit the work for publication.

We thank Christina Aaby Svendsen (Technical University of Denmark) for technical support with the Illumina sequencing of the plasmidomes.

Sequencing data analysis was performed using the DeiC National Life Science Supercomputer at DTU.

## Data availability

The DNA sequences generated in this project are available through ENA/GenBank/DDBJ under the accession number PRJEB41171 (Nanopore reads: ERX4715074-ERX4715097; Illumina reads: ERX5299122-ERX5299145; Assemblies: ERZ1694234-ERZ1694257). The code for the creation of assemblies is accessible from Github (https://github.com/philDTU/plasmidPublication) and additional supplementary material is available at https://figshare.com/projects/A_Peek_into_the_Plasmidome_of_Global_Sewage/94448.

## Figure legends

**Table S1**. Sewage sample information.

**Figure S1: Length of nanopore sequencing reads**. The violin plot displays log transformed read lengths. The horizontal dashed lines indicate log values for 1.000 and 10.000 bases length, respectively. Most reads exhibit a read length below 10.000 bases, which is the cut-off value for our assembly workflow, and most of the reads are between 1.000 and 10.000 bases long.

**Figure S2. Plasmid and virus (phage)-related circular DNA elements**. The bar plots display the fraction (A) and total counts (B) of circular contigs containing Pfam IDs specific for plasmid and virus/phage -related proteins per sample. Each predicted protein by prodigal was searched against the pfam databases using HMMER hmmscan and filtered for a p-value less than 0.00001. In a small subset of assemblies we identify both viral and plasmid associated genes. Pfam ID’s classified as “other than plasmid & viral“ might still be plasmid relevant; they are just not specified as plasmid-related based on the stringent scheme used.

**Figure S3. Comparison to known plasmids in plasmid database (PLSDB) – clustering on individual plasmid level**. Samples with less than 100 circular assembled contigs where remove from the analysis as well as plasmids with less than 10 occurrences over all samples. Clustering of samples (columns) was done using Euclidean distance of the centered log ratio (clr)-transformed values.

**Figure S4. Comparison between plasmidome samples – MASH distances**. All plasmid candidate sequences for each sample from the five examined continents were sketched using MASH, distances calculated, and visualized by principal component analysis. A) This plot displays the differences and similarities between all 24 plasmidome samples. B) This plot displays the differences and similarities between 22 plasmidome samples (all samples, except NGA and BRA.2).

**Figure S5. Heatmaps depicting antimicrobial resistance profiles from the whole community and plasmidomes from global sewage** based on presence/absence (A) and centered log ratio (clr)-transformed abundancies (B) of antimicrobial resistance gene classes. The antimicrobial resistance genes were identified in a ResFinder-based analysis using the Illumina reads from the whole community, Illumina reads from the plasmid preparations, and Nanopore reads from the plasmid preparations.

**Figure S6. Functional characterization of circular DNA elements – GO annotation**. The heat map displays centered log ratio (clr)-transformed abundancies of GO annotations assigned to predicted proteins. Samples with less than 100 circular assembled contig were remove from the analysis as well as GO identifiers with less than 10 occurrences over all samples. The clustering of samples was performed using Euclidean distance of the clr-transformed values resulting in 2 main clusters.

**Figure S7. Comparison of AMR genes with prevalence data by CARD (https://card.mcmaster.ca)**. The most frequently observed AMR genes that were more abundant in plasmidomes (as compared to in the whole community sequencing data) were explored at the CARD website. Here, the prevalence for AMR genes is presented for a selection of pathogens, whether they are associated with the plasmid or chromosome. The prevalence data are calculated as follows: Antimicrobial resistance (AMR) molecular prevalence data were generated using the Resistance Gene Identifier (RGI), a tool for putative AMR gene detection from submitted sequence data using the AMR detection models available in CARD. To generate prevalence data, RGI was used to analyze molecular sequence data available in NCBI Genomes for 88 pathogens of interest. For each of these pathogens, complete chromosome sequences, complete plasmid sequences, and whole genome shotgun (WGS) assemblies were analyzed individually by RGI. RGI results were then aggregated to calculate percent occurrence. (See also Alcock *et al*., NAR, 2020, https://academic.oup.com/nar/article/48/D1/D517/5608993).

## Notes

https://figshare.com/projects/A_Peek_into_the_Plasmidome_of_Global_Sewage/94448

### Competing Interest Statement

The authors have declared no competing interest.

## References

1. Lederberg J. 1952. Cell Genetics and Hereditary Symbiosis. Physiological Reviews 32:403–430.

2. Cohen SN, Chang ACY, Boyer HW, Helling RB. 1973. Construction of Biologically Functional Bacterial Plasmids In Vitro. PNAS 70:3240–3244.

3. Rodríguez-Beltrán J, DelaFuente J, León-Sampedro R, MacLean RC, San Millán Á. 2021. Beyond horizontal gene transfer: the role of plasmids in bacterial evolution. Nature Reviews Microbiology https://doi.org/10.1038/s41579-020-00497-1.

4. Johnson TJ, Nolan LK. 2009. Pathogenomics of the Virulence Plasmids of Escherichia coli. Microbiol Mol Biol Rev 73:750–774.

5. Bratu S, Brooks S, Burney S, Kochar S, Gupta J, Landman D, Quale J. 2007. Detection and Spread of Escherichia coli Possessing the Plasmid-Borne Carbapenemase KPC-2 in Brooklyn, New York. Clin Infect Dis 44:972–975.

6. Tian G-B, Doi Y, Shen J, Walsh TR, Wang Y, Zhang R, Huang X. 2017. MCR-1-producing Klebsiella pneumoniae outbreak in China. The Lancet Infectious Diseases 17:577.

7. Hendriksen RS, Munk P, Njage P, Bunnik B, McNally L, Lukjancenko O, Röder T, Nieuwenhuijse D, Pedersen SK, Kjeldgaard J, Kaas RS, Clausen PTLC, Vogt JK, Leekitcharoenphon P, Schans MGM, Zuidema T, Husman AMR, Rasmussen S, Petersen B, Bego A, Rees C, Cassar S, Coventry K, Collignon P, Allerberger F, Rahube TO, Oliveira G, Ivanov I, Vuthy Y, Sopheak T, Yost CK, Ke C, Zheng H, Baisheng L, Jiao X, Donado-Godoy P, Coulibaly KJ, Jergović M, Hrenovic J, Karpíšková R, Villacis JE, Legesse M, Eguale T, Heikinheimo A, Malania L, Nitsche A, Brinkmann A, Saba CKS, Kocsis B, Solymosi N, Thorsteinsdottir TR, Hatha AM, Alebouyeh M, Morris D, Cormican M, O’Connor L, Moran-Gilad J, Alba aP, Battisti A, Shakenova Z, Kiiyukia C, Ng’eno E, Raka L, Avsejenko J, Bērziņš A, Bartkevics V, Penny C, Rajandas H, Parimannan S, Haber MV, Pal P, Jeunen G-J, Gemmell N, Fashae K, Holmstad R, Hasan R, Shakoor S, Rojas MLZ, Wasyl D, Bosevska G, Kochubovski M, Radu C, Gassama A, Radosavljevic V, Wuertz S, Zuniga-Montanez R, Tay MYF, Gavačová D, Pastuchova K, Truska P, Trkov M, Esterhuyse K, Keddy K, Cerdà-Cuéllar M, Pathirage S, Norrgren L, Örn S, Larsson DGJ, Van der Heijden T, Kumburu HH, Sanneh B, Bidjada P, Njanpop-Lafourcade B-M, Nikiema-Pessinaba SC, Levent B, Meschke JS, Beck NK, Van CD, Do Phuc N, Tran DMN, Kwenda G, Tabo D, Wester AL, Cuadros-Orellana S, Amid C, Cochrane G, Sicheritz-Ponten T, Schmitt H, Alvarez JRM, Aidara-Kane A, Pamp SJ, Lund O, Hald T, Woolhouse M, Koopmans MP, Vigre H, Petersen TN, Aarestrup FM. 2019. Global monitoring of antimicrobial resistance based on metagenomics analyses of urban sewage. Nature Communications 10:1124.

8. Munk P, Knudsen BE x000E6 ER, Lukjacenko O, Duarte ASR, Gompel L, Luiken REC, Smit LAM, Schmitt H, Garcia AD, Hansen RB, Petersen TN, Bossers A, x000E9 ER, Graveland H, van Essen A, Gonzalez-Zorn B, Moyano G, Sanders P, Chauvin C, David J, Battisti A, Caprioli A, Dewulf J, Blaha T, Wadepohl K, Brandt M, Wasyl D, ska MS x00144, Zajac M, Daskalov H, Saatkamp HW, rk KDCS x000E4, Lund O, Hald T, Pamp S x000FC nje, Vigre H x000E5 kan, Heederik D, Wagenaar JA, Mevius D, Aarestrup FM. 2018. Abundance and diversity of the faecal resistome in slaughter pigs and broilers in nine European countries. Nature Microbiology 1–14.

9. Campbell TP, Sun X, Patel VH, Sanz C, Morgan D, Dantas G. 2020. The microbiome and resistome of chimpanzees, gorillas, and humans across host lifestyle and geography. 6. The ISME Journal 14:1584–1599.

10. Chen Q-L, Cui H-L, Su J-Q, Penuelas J, Zhu Y-G. 2019. Antibiotic Resistomes in Plant Microbiomes. Trends in Plant Science 24:530–541.

11. Forsberg KJ, Patel S, Gibson MK, Lauber CL, Knight R, Fierer N, Dantas G. 2014. Bacterial phylogeny structures soil resistomes across habitats. 7502. Nature 509:612–616.

12. Carr VR, Witherden EA, Lee S, Shoaie S, Mullany P, Proctor GB, Gomez-Cabrero D, Moyes DL. 2020. Abundance and diversity of resistomes differ between healthy human oral cavities and gut. 1. Nature Communications 11:693.

13. Wick RR, Judd LM, Gorrie CL, Holt KE. 2017. Unicycler: Resolving bacterial genome assemblies from short and long sequencing reads. PLOS Computational Biology 13:e1005595.

14. Antipov D, Hartwick N, Shen M, Raiko M, Lapidus A, Pevzner PA. 2016. plasmidSPAdes: assembling plasmids from whole genome sequencing data. Bioinformatics 32:3380–3387.

15. Vielva L, de Toro M, Lanza VF, de la Cruz F. 2017. PLACNETw: a web-based tool for plasmid reconstruction from bacterial genomes. Bioinformatics 33:3796–3798.

16. Rozov R, Brown Kav A, Bogumil D, Shterzer N, Halperin E, Mizrahi I, Shamir R. 2017. Recycler: an algorithm for detecting plasmids from de novo assembly graphs. Bioinformatics 33:475–482.

17. Dean FB, Nelson JR, Giesler TL, Lasken RS. 2001. Rapid Amplification of Plasmid and Phage DNA Using Phi29 DNA Polymerase and Multiply-Primed Rolling Circle Amplification. Genome Res 11:1095–1099.

18. Che Y, Xia Y, Liu L, Li A-D, Yang Y, Zhang T. 2019. Mobile antibiotic resistome in wastewater treatment plants revealed by Nanopore metagenomic sequencing 1–13.

19. Bertrand D, Shaw J, Kalathiyappan M, Ng AHQ, Kumar MS, Li C, Dvornicic M, Soldo JP, Koh JY, Tong C, Ng OT, Barkham T, Young B, Marimuthu K, Chng KR, Sikic M, Nagarajan N. 2019. Hybrid metagenomic assembly enables high-resolution analysis of resistance determinants and mobile elements in human microbiomes. Nature Biotechnology 1–15.

20. Antipov D, Raiko M, Lapidus A, Pevzner PA. 2019. Plasmid detection and assembly in genomic and metagenomic datasets. Genome Res gr. 241299.118.

21. Jørgensen TS, Hansen MA, Xu Z, Tabak MA, Sørensen SJ, Hansen LH. 2017. Plasmids, Viruses, And Other Circular Elements In Rat Gut. bioRxiv 143420.

22. Kav AB, Sasson G, Jami E, Doron-Faigenboim A, Benhar I, Mizrahi I. 2012. Insights into the bovine rumen plasmidome. PNAS 109:5452–5457.

23. Kav AB, Rozov R, Bogumil D, Sørensen SJ, Hansen LH, Benhar I, Halperin E, Shamir R, Mizrahi I. 2020. Unravelling plasmidome distribution and interaction with its hosting microbiome. Environmental Microbiology 22:32–44.

24. Zhang T, Zhang X-X, Ye L. 2011. Plasmid Metagenome Reveals High Levels of Antibiotic Resistance Genes and Mobile Genetic Elements in Activated Sludge. PLOS ONE 6:e26041.

25. Sentchilo V, Mayer AP, Guy L, Miyazaki R, Green Tringe S, Barry K, Malfatti S, Goessmann A, Robinson-Rechavi M, van der Meer JR. 2013. Community-wide plasmid gene mobilization and selection. 6. The ISME Journal 7:1173–1186.

26. Kothari A, Wu Y-W, Chandonia J-M, Charrier M, Rajeev L, Rocha AM, Joyner DC, Hazen TC, Singer SW, Mukhopadhyay A. 2019. Large Circular Plasmids from Groundwater Plasmidomes Span Multiple Incompatibility Groups and Are Enriched in Multimetal Resistance Genes. mBio 10.

27. Kav AB, Sasson G, Jami E, Doron-Faigenboim A, Benhar I, Mizrahi I. 2012. Insights into the bovine rumen plasmidome. PNAS 109:5452–5457.

28. Li H. 2018. Minimap2: pairwise alignment for nucleotide sequences. Bioinformatics 34:3094–3100.

29. Li H. 2016. Minimap and miniasm: fast mapping and de novo assembly for noisy long sequences. Bioinformatics 32:2103–2110.

30. Vaser R, Sović I, Nagarajan N, šikić M. 2017. Fast and accurate de novo genome assembly from long uncorrected reads. Genome Res 27:737–746.

31. Walker BJ, Abeel T, Shea T, Priest M, Abouelliel A, Sakthikumar S, Cuomo CA, Zeng Q, Wortman J, Young SK, Earl AM. 2014. Pilon: An Integrated Tool for Comprehensive Microbial Variant Detection and Genome Assembly Improvement. PLOS ONE 9:e112963.

32. Galata V, Fehlmann T, Backes C, Keller A. 2019. PLSDB: a resource of complete bacterial plasmids. Nucleic Acids Res 47:D195–D202.

33. Ondov BD, Treangen TJ, Melsted P, Mallonee AB, Bergman NH, Koren S, Phillippy AM. 2016. Mash: fast genome and metagenome distance estimation using MinHash. Genome Biology 17:132.

34. Zankari E, Hasman H, Cosentino S, Vestergaard M, Rasmussen S, Lund O, Aarestrup FM, Larsen MV. 2012. Identification of acquired antimicrobial resistance genes. Journal of Antimicrobial Chemotherapy 67:2640–2644.

35. Clausen PTLC, Aarestrup FM, Lund O. 2018. Rapid and precise alignment of raw reads against redundant databases with KMA. BMC Bioinformatics 19:307.

36. Hyatt D, Chen G-L, Locascio PF, Land ML, Larimer FW, Hauser LJ. 2010. Prodigal: prokaryotic gene recognition and translation initiation site identification. BMC Bioinformatics 11:11:119.

37. El-Gebali S, Mistry J, Bateman A, Eddy SR, Luciani A, Potter SC, Qureshi M, Richardson LJ, Salazar GA, Smart A, Sonnhammer ELL, Hirsh L, Paladin L, Piovesan D, Tosatto SCE, Finn RD. 2019. The Pfam protein families database in 2019. Nucleic Acids Res 47:D427–D432.

38. Mitchell A, Chang H-Y, Daugherty L, Fraser M, Hunter S, Lopez R, McAnulla C, McMenamin C, Nuka G, Pesseat S, Sangrador-Vegas A, Scheremetjew M, Rato C, Yong S-Y, Bateman A, Punta M, Attwood TK, Sigrist CJA, Redaschi N, Rivoire C, Xenarios I, Kahn D, Guyot D, Bork P, Letunic I, Gough J, Oates M, Haft D, Huang H, Natale DA, Wu CH, Orengo C, Sillitoe I, Mi H, Thomas PD, Finn RD. 2015. The InterPro protein families database: the classification resource after 15 years. Nucleic Acids Res 43:D213–D221.

39. Jørgensen TS, Hansen MA, Xu Z, Tabak MA, Sørensen SJ, Hansen LH. 2017. Plasmids, Viruses, And Other Circular Elements In Rat Gut. bioRxiv 143420.

40. Katoh K, Standley DM. 2013. MAFFT Multiple Sequence Alignment Software Version 7: Improvements in Performance and Usability. Molecular Biology and Evolution 30:772–780.

41. Afgan E, Baker D, Batut B, van den Beek M, Bouvier D, Čech M, Chilton J, Clements D, Coraor N, Grüning BA, Guerler A, Hillman-Jackson J, Hiltemann S, Jalili V, Rasche H, Soranzo N, Goecks J, Taylor J, Nekrutenko A, Blankenberg D. 2018. The Galaxy platform for accessible, reproducible and collaborative biomedical analyses: 2018 update. Nucleic Acids Research 46:W537–W544.

42. Price MN, Dehal PS, Arkin AP. 2010. FastTree 2 – Approximately Maximum-Likelihood Trees for Large Alignments. PLOS ONE 5:e9490.

43. Seemann T. 2014. Prokka: rapid prokaryotic genome annotation. Bioinformatics 30:2068–2069.

44. Carver T, Thomson N, Bleasby A, Berriman M, Parkhill J. 2009. DNAPlotter: circular and linear interactive genome visualization. Bioinformatics 25:119–120.

45. Altschul SF, Gish W, Miller W, Myers EW, Lipman DJ. 1990. Basic local alignment search tool. Journal of molecular biology 215:403–410.

46. Lanciano S, Carpentier M-C, Llauro C, Jobet E, Robakowska-Hyzorek D, Lasserre E, Ghesquière A, Panaud O, Mirouze M. 2017. Sequencing the extrachromosomal circular mobilome reveals retrotransposon activity in plants. PLOS Genetics 13:e1006630.

47. Shibata Y, Kumar P, Layer R, Willcox S, Gagan JR, Griffith JD, Dutta A. 2012. Extrachromosomal microDNAs and chromosomal microdeletions in normal tissues. Science 336:82–86.

48. Møller HD, Mohiyuddin M, Prada-Luengo I, Sailani MR, Halling JF, Plomgaard P, Maretty L, Hansen AJ, Snyder MP, Pilegaard H, Lam HYK, Regenberg B. 2018. Circular DNA elements of chromosomal origin are common in healthy human somatic tissue. 1. Nature Communications 9:1069.

49. San Millan A. 2018. Evolution of Plasmid-Mediated Antibiotic Resistance in the Clinical Context. Trends in Microbiology 26:978–985.

50. Ogilvie LA, Firouzmand S, Jones BV. 2012. Evolutionary, ecological and biotechnological perspectives on plasmids resident in the human gut mobile metagenome. Bioengineered 3:13–31.

51. Alcock BP, Raphenya AR, Lau TTY, Tsang KK, Bouchard M, Edalatmand A, Huynh W, Nguyen A-LV, Cheng AA, Liu S, Min SY, Miroshnichenko A, Tran H-K, Werfalli RE, Nasir JA, Oloni M, Speicher DJ, Florescu A, Singh B, Faltyn M, Hernandez-Koutoucheva A, Sharma AN, Bordeleau E, Pawlowski AC, Zubyk HL, Dooley D, Griffiths E, Maguire F, Winsor GL, Beiko RG, Brinkman FSL, Hsiao WWL, Domselaar GV, McArthur AG. 2020. CARD 2020: antibiotic resistome surveillance with the comprehensive antibiotic resistance database. Nucleic Acids Res 48:D517–D525.

52. Stalder T, Press MO, Sullivan S, Liachko I, Top EM. 2019. Linking the resistome and plasmidome to the microbiome. 10. The ISME Journal 13:2437–2446.

53. Lan F, Demaree B, Ahmed N, Abate AR. 2017. Single-cell genome sequencing at ultra-high-throughput with microfluidic droplet barcoding. Nature Biotechnology 35:640–646.

