## Supplementary material for "A Peek into the Plasmidome of Global Sewage": Table S1

| sample | continent | Country | city | extraction_date | plasmid_safe_dnase_date | phi20_date | sequencing_date | run_id |
|---|---|---|---|---|---|---|---|---|
| Africa - CIV | Africa | Côte d'Ivoire | Abidjan | 25/03/2019 | 03/04/2019 | 04/04/2019 | 10/04/2019 | GS_Plasmidome_13 |
| Africa - GHA | Africa | Ghana | Tamale | 25/03/2019 | 03/04/2019 | 04/04/2019 | 05/04/2019 | GS_Plasmidome_4 |
| Africa - NGA | Africa | Nigeria | NA | 28/03/2019 | 15/04/2019 | 16/04/2019 | 06/05/2019 | GS_Plasmidome_50 |
| Africa - TZA | Africa | Tanzania | Moshi | 01/04/2019 | 15/04/2019 | 16/04/2019 | 16/05/2019 | GS_Plasmidome_15 |
| Asia - CHN | Asia | China | Guangzhou | 01/04/2019 | 15/04/2019 | 16/04/2019 | 13/05/2019 | GS_Plasmidome_64 |
| Asia - IND | Asia | India | Cochin | 25/03/2019 | 03/04/2019 | 04/04/2019 | 05/04/2019 | GS_Plasmidome_11 |
| Asia - KHM | Asia | Cambodia | Phnom Penh | 26/03/2019 | 03/04/2019 | 04/04/2019 | 12/04/2019 | GS_Plasmidome_21 |
| Asia - NPL | Asia | Nepal | Kathmandu | 27/03/2019 | 15/04/2019 | 16/04/2019 | 16/05/2019 | GS_Plasmidome_33 |
| Asia - PAK | Asia | Pakistan | Karachi | 25/03/2019 | 03/04/2019 | 04/04/2019 | 08/04/2019 | GS_Plasmidome_7 |
| Europe - ALB | Europe | Albania | Tirana | 26/03/2019 | 03/04/2019 | 04/04/2019 | 10/04/2019 | GS_Plasmidome_17 |
| Europe - CZE | Europe | Czechia | Brno | 27/03/2019 | 03/04/2019 | 04/04/2019 | 29/04/2019 | GS_Plasmidome_23 |
| Europe - DEU | Europe | Germany | Berlin | 27/03/2019 | 03/04/2019 | 04/04/2019 | 29/04/2019 | GS_Plasmidome_27 |
| Europe - ESP | Europe | Spain | Barcelona | 01/04/2019 | 15/04/2019 | 16/04/2019 | 20/05/2019 | GS_Plasmidome_75 |
| Europe - ISL | Europe | Iceland | Reykjavik | 27/03/2019 | 03/04/2019 | 04/04/2019 | 01/05/2019 | GS_Plasmidome_28 |
| Europe - POL | Europe | Poland | Pu_awy | 28/03/2019 | 15/04/2019 | 16/04/2019 | 03/05/2019 | GS_Plasmidome_36 |
| Europe - SVN | Europe | Slovenia | Ljubljana | 28/03/2019 | 15/04/2019 | 16/04/2019 | 03/05/2019 | GS_Plasmidome_38 |
| Europe - XK | Europe | Kosovo | Prishtina | 01/04/2019 | 15/04/2019 | 16/04/2019 | 08/05/2019 | GS_Plasmidome_60 |
| North America - CAN | North America | Canada | Toronto | 26/03/2019 | 03/04/2019 | 04/04/2019 | 12/04/2019 | GS_Plasmidome_22b |
| North America - USA.1 | North America | United States of America | Portland,OR | 01/04/2019 | 15/04/2019 | 16/04/2019 | 13/05/2019 | GS_Plasmidome_74d |
| North America - USA.2 | North America | United States of America | El Paso, Texas | 01/04/2019 | 15/04/2019 | 16/04/2019 | 20/05/2019 | GS_Plasmidome_74e |
| South America - BRA.1 | South America | Brasil | Belo Horizonte | 28/03/2019 | 15/04/2019 | 16/04/2019 | 08/05/2019 | GS_Plasmidome_53 |
| South America - BRA.2 | South America | Brasil | Belém | 28/03/2019 | 15/04/2019 | 16/04/2019 | 06/05/2019 | GS_Plasmidome_53a |
| South America - ECU | South America | Ecuador | Galapagos | 25/03/2019 | 03/04/2019 | 04/04/2019 | 08/04/2019 | GS_Plasmidome_14a |
| South America - PER | South America | Peru | Lima | 27/03/2019 | 15/04/2019 | 16/04/2019 | 01/05/2019 | GS_Plasmidome_35 |
